# Tagging fluorescent reporter to epinecidin-1 antimicrobial peptide

**DOI:** 10.1101/2025.06.12.659415

**Authors:** Sivakumar Jeyarajan, Anbarasu Kumarasamy

## Abstract

In this study, we successfully cloned the fluorescent proteins eGFP and DsRed in-frame with the antimicrobial peptide epinecidin-1 (FIFHIIKGLFHAGKMIHGLV). The cloning strategy involved inserting the fluorescent reporters into the expression vector, followed by screening of positive clones through visual fluorescence detection and molecular validation. The visually identified fluorescent colonies were found positive with PCR and plasmid migration assay confirming successful cloning. This fusion of fluorescent reporters with a short antimicrobial peptide enables real-time visualization and monitoring of the peptide’s mechanism of action on membranes and within cells, both in vivo and in vitro. The fusion of eGFP and DsRed to epinecidin-1 did not impair the expression or fluorescence of the reporter protein.

## 1. Introduction

Genetically encoded tags, such as fluorescent proteins, are powerful tools for studying the dynamics of proteins and peptides using fluorescence microscopy or other fluorescence-based detection systems. The incorporation of a fluorescent tag as a reporter enables precise localization of individual protein or peptide molecules within the microscopic resolution of cells. One of the most transformative developments in this field has been the discovery of green fluorescent protein (GFP), originally derived from the jellyfish *Aequorea victoria* [1]. GFP has revolutionized biological imaging and become an essential tool for observing cellular processes *in vivo* [2, 3]. Since its introduction, extensive protein engineering has led to the development of a wide array of fluorescent proteins with diverse colour palettes and enhanced properties [4]. These diverse spectrum of palettes derived from GFP exhibit significant variation in their spectral characteristics, allowing for specialized applications in biological imaging [5]. The use of fluorescent proteins, particularly GFP and DsRed from *Discosoma striata*, has enabled researchers to quantitatively analyze protein diffusion, exchange, and targeted localization in living cells, offering unprecedented insights into cellular dynamics [6].

Green Fluorescent Protein (GFP), composed of 238 amino acids, adopts a compact tertiary structure characterized by six alpha helices and eleven beta strands arranged in a β-barrel configuration, interconnected by flexible loops. The intrinsic fluorescence of GFP originates from a chromophore formed through the post-translational cyclization of a serine-tyrosine-glycine tripeptide, followed by oxidation of the tyrosine residue. Importantly, this process occurs autonomously, without the requirement for exogenous cofactors, substrates, or enzymatic activity, thereby facilitating its application in live-cell imaging [6]. GFP fusion constructs are typically generated by attaching the protein of interest to either at the N- or C-terminus of GFP. Given the spatial proximity of these termini, they can be linked *via* short peptide sequences, preserving the structural integrity and fluorescence of the fusion protein. The exceptional stability of GFP under a wide range of chemical and physical conditions has further contributed to its widespread use as a molecular marker [7] [8] particularly in golgi apparatus and lysosomes. The versatility of GFP has enabled real-time visualization of numerous cellular and developmental processes *in vivo*. These include, but are not limited to, gene expression profiling, sub-cellular protein localization, protein-protein interactions, cell cycle progression, chromosomal dynamics, intracellular trafficking, and organelle biogenesis [9, 10]. Moreover, GFP can function as a reporter gene under the control of specific promoters, with fluorescence intensity serving as a direct proxy for transcriptional activity in living cells and tissues [11]. A significant advancement in GFP technology was the development of enhanced GFP (eGFP) through codon optimization, which improved expression efficiency and fluorescence intensity in a broad range of transgenic organisms [12]. These innovations have solidified GFP as an indispensable tool in molecular and cellular biology.

Antimicrobial peptides (AMPs) represent a structurally diverse and functionally versatile class of molecules, typically ranging from 12 to 50 amino acids in length. They are categorized into subgroups based on their amino acid composition and structural motifs [13]. The secondary structures of AMPs generally conform to one of four major types: (i) α-helical structures, (ii) β-stranded conformations stabilized by two or more disulfide bonds, (iii) β-hairpin or loop structures formed by a single disulfide bond and/or peptide cyclization, and (iv) extended conformations. Many AMPs are intrinsically unstructured in aqueous solution but undergo conformational folding upon interaction with biological membranes. This membrane-induced folding often results in amphipathic structures, where hydrophilic residues align along one face of the helix and hydrophobic residues along the opposite face, facilitating membrane association and disruption.

Antimicrobial peptides (AMPs) are evolutionarily conserved components of the innate immune system, present across a wide range of organisms including bacteria, plants, invertebrates, and vertebrates. Also referred to as peptide antibiotics, AMPs exhibit broad-spectrum antimicrobial activity and are less prone to inducing resistance compared to conventional antibiotics. Remarkably, many bacterial species have retained sensitivity to AMPs despite extensive evolutionary pressure. Unlike the adaptive immune system, which relies on immunological memory to mount specific and enhanced responses upon re-exposure to pathogens, AMPs function through a distinct, non-specific mechanism of action. Most AMPs are effective against a wide array of pathogens, making them particularly valuable for treating both localized and systemic infections[14, 15]. Beyond their antimicrobial properties, several AMPs also exhibit immunomodulatory functions. For instance, defensins have been implicated in the recruitment of effector T cells to sites of inflammation, thereby contributing to the effector phase of adaptive immunity [16]. These multifunctional characteristics make AMPs as promising candidates for the development of novel therapeutic agents.

In this study, epinecidin-1 an antimicrobial peptide originally identified in groupers (*Epinephelus coioides*) which was reported to have antibacterial activity against both Gram-negative and Gram-positive bacteria [17-19] and antifungal properties [20, 21] [22] was cloned in-frame with eGFP and DsRed.

## 2. Materials and Methods

### 2.1. Bacterial Strains, Vectors, and Reagents

The *Escherichia coli* (*E. coli*) DH5α strain was used for cloning. Restriction enzymes (BamHI, EcoRI, XhoI) and T4 DNA ligase were obtained from New England Biolabs (Ipswich, MA). The plasmid vector pET-32a-Epi, previously described in [19], was used for cloning the GFP and DsRed genes upstream of the antimicrobial peptide Epi-1. Luria-Bertani (LB) broth and agar were purchased from Sigma-Aldrich.

### 2.2 Preparation of Competent Cells and Transformation

Competent *E. coli* DH5α cells were prepared by growing a few colonies in 100 mL of LB broth to mid-log phase, followed by chilling on ice for 15 minutes. Cells were harvested by centrifugation at 4,000 rpm for 10 minutes at 4°C, washed with 30 mL of ice-cold divalent cation solution (80 mM MgCl_2_, 20 mM CaCl_2_), and centrifuged again.

The pellet was then washed with 10 mL of 100 mM CaCl_2_, centrifuged, and finally resuspended in 1 mL of 100 mM CaCl_2_. For transformation, 100 µL of competent cells were mixed with 10 ng of plasmid DNA, incubated on ice for 30 minutes, heat-shocked at 42°C for 90 seconds, and returned to ice for 10 minutes. After recovery, 800 µL LB medium was added and incubated at 37°C for 1 hour, cells were plated on LB agar containing 100 µg/mL ampicillin and incubated overnight at 37°C in an inverted position.

### 2.3. Plasmid isolation

A single transformed colony was inoculated into 10 mL LB broth with 100 µg/mL ampicillin and incubated overnight at 37°C with shaking. Cells were pelleted by centrifugation at 10,000 rpm for 10 minutes at 4°C. The pellet was resuspended in 250 µL of Solution I (50 mM glucose, 25 mM Tris-HCl pH 8.0, 10 mM EDTA), followed by the addition of 250 µL of Solution II (0.2 N NaOH, 1% SDS). After gentle mixing, 350 µL of Solution III (5 mM potassium acetate with glacial acetic acid) was added. The lysate was centrifuged at 10,000 rpm for 5 minutes at 4°C, and the supernatant was transferred to a fresh tube. Plasmid DNA was precipitated with 0.6 volumes of isopropanol, centrifuged, washed with 70% ethanol, air-dried, and resuspended in 50 µL of double-distilled water. DNA concentration was measured spectrophotometrically and stored at 4°C (short-term) or −20°C (long-term). Plasmids used included pET32a-Epi-1, pEGFP-C1 (courtesy of Dr. J.Y. Chen, Academia Sinica, Taiwan), and pDsRed-C1 Monomer (Clontech).

### 2.4. PCR Amplification of eGFP and DsRed

Target genes were amplified using the following primers: For eGFP amplification, forward (BamHI): 5′-GGCGTGGATCCATGGTGAGCAAGGGCGAGGAG-3′ and reverse (EcoRI): 5′-GATCCGAATTCCTTGTACAGCTCGTCCATGCCG-3′ primers were used. For DsRed amplification, forward (BamHI): 5′-CCATGGATCCATGGACAACACCGAG-3′ and reverse (EcoRI): 5′-GGTGGAATTCCTGGGAGCCGGAG-3′ primers were used. The restriction sites in the primers are italicized and underlined. PCR was performed using Taq polymerase (Takara, Japan). Products were resolved on 1.2% TAE agarose gel, visualized under UV light, and extracted using the QIAquick Gel Extraction Kit (QIAGEN, USA).

### 2.5. Restriction digestion

PCR products and the pET-32a-Epi vector were digested with BamHI and EcoRI (NEB) at 37°C for 4 hours. Digested products were resolved on 1% agarose gel, and the desired bands were excised and purified using the QIAquick Gel Extraction Kit.

### 2.6. Ligation and Transformation

Ligation reactions (10 µL) were set up using the digested vector and either eGFP or DsRed insert, incubated overnight at 16°C. Five microliters of the ligation mix was used to transform 100 µL of competent *E. coli* DH5α cells as described in Section 2.2

### 2.7. Colony Patching and Screening

Transformed colonies were individually picked using sterile toothpicks and patched onto LB agar plates containing ampicillin. Colonies were arranged in a grid format and screened for the presence of inserts using colony PCR with gene-specific primers.

### 2.8. Boiling Lysis and Colony PCR

For preliminary screening, NF (Non-Fluorescent) and F (Fluorescent) colonies were dissolved in 15 µL of sterile deionized water, boiled at 94°C for 10 minutes, and centrifuged at 14,000 rpm for 10 minutes. The supernatant was used as the PCR template for amplification with eGFP or DsRed-specific primers.

### 2.9. Colony Patching and Screening with insert specific and spanning PCR

Transformed colonies were individually picked using sterile toothpicks and patched onto LB agar plates containing ampicillin. Colonies were arranged in a grid format and screened for the presence of inserts using insert specific primers using eGFP or DsRed specific primers (amplicon length 717 bp for eGFP and 685 bp for DsRed) and spanning PCR using fluorescent protein forward primer and Epi-1 reverse primer to confirm the correct orientation of fluorescent reporter in frame with Epi-1. The size of the amplicon for spanning PCR are eGFP-epi-1 (717+84=801bp) and Ds-Red-Epi-1 is (685+84=769bp). Plasmid was isolated from colonies as described in 2..3

## 3. Results

### 3.1. Construction of fluorescent Epi-1 vector

In our previous studies, we cloned the Epi-1 gene into the pET32a vector for antimicrobial peptide over expression. To that vector, eGFP and DsRed genes were inserted between BamH1 and EcoRI sites. The schematic of the constructs are shown in Figure-1. The genes are driven by T_7_ promoter to express in the bacterial host. The start codon is in the thioredoxin protein, followed by His-tag for purification, followed by enterokinase site for cleavage. The fluorescent reporter eGFP or DsRed is after the enterokinase cleavage site. The thioredoxin helps in protein expression based on its chaperone properties. The thioredoxin protein can be removed by cleaving with enterokinase and captured by his tag binding resin releasing the fluorescent tagged Epi-1. The advantage of this vector is the thioredoxin gene if not needed, it can be removed by NdeI enzyme digestion [23] so only the GFP and antimicrobial peptide gene will be in frame and can be purified by His-Tag affinity chromatography. After the active epi-1 gene, a stop codon is present. Following the stop codon is the T_7_-terminator signal to terminate the transcription. The size of pET32a-Epi-1 vector before fluorescent reporter gene ligation is 5944 bp. After ligation of eGFP insert (717 bp) and DsRed insert (685 bp), the resultant pET32a-eGFP-epi clone is 6661 bp and pET32a-DsRed-epi-1 clone is 6629 bp. As shown in the figure, The Epi-1 following the fluorescent protein is shiwn in pink. The expression cassette contains a stop codon after Epi-1 gene followed by T7 termination signal to stop transcript synyhesis after compeltion of the required portion synthesized.

### 3.2. Visual screen of fluorescent reporter -Epi-1 clones

After the molecular cloning strategies were designed for incorporating the fusion fluorescent reporter genes to Epi-1, pET32a-Epi-1 vector was digested using the restriction enzymes BamHI and *Eco*RI, resulting in a single linear DNA band. The eGFP and DsRed inserts were PCR-amplified from pEGFP-C1 and pDsRed-Monomer from Clonetech. The amplified products were also digested with the same enzyme pair of restriction enzymes. The ligation of inserts to pET32a-Epi-1 vector yielded numerous colonies after spreading on the LB-agar plate. After 48 hours, the fluorescence in colonies were observed. However the insert positive colonies did not show any fluorescence after 24 hours. As shown in Figure 2, the enlarged plate photographs show a clear distinction of fluorescent and non-fluorescent colonies. The fluorescent colonies were subjected to PCR and plasmid isolation. Simultaneously the non-fluorescent colonies were used for molecular charcterization as controls.

**Figure 1.**
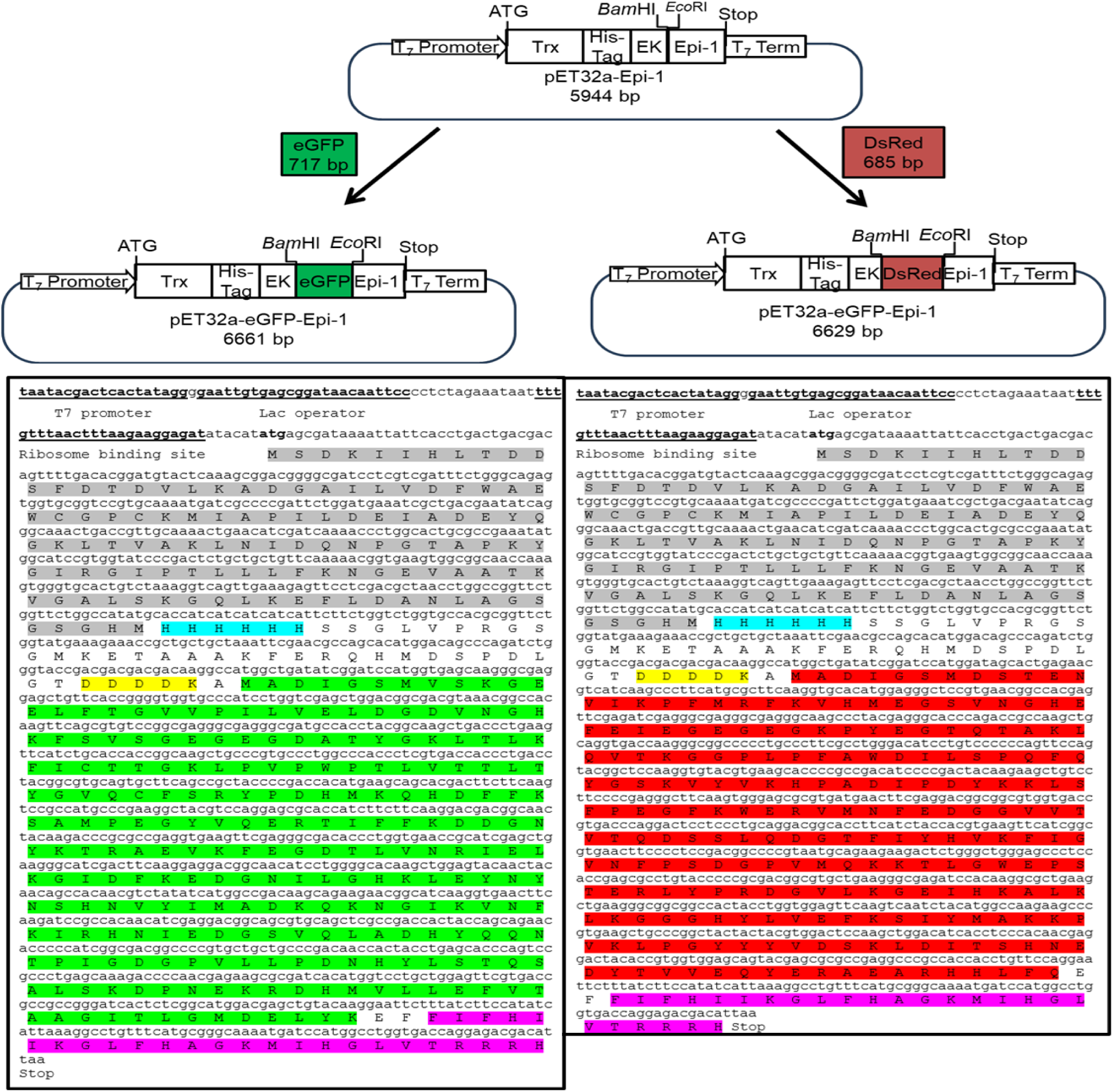
Plasmid Constructs for Fluorescent Epi-1 Expression. This figure presents annotated maps and DNA sequences of two recombinant plasmids: pET32a-GFP-Epi-1 and pET32a-DsRed-Epi-1. The eGFP (717 bp) and DsRed (685 bp) genes were individually inserted into the pET32a-Epi-1 vector (5944 bp), resulting in total plasmid sizes of 6661 bp for pET32a-GFP-Epi-1 and 6629 bp for pET32a-DsRed-Epi-1, respectively. The constructs feature key regulatory and expression elements, including a T7 promoter, lac operator, ribosome binding site, Trx tag, His-tag, enterokinase cleavage site, and the Epi-1 gene. The DNA sequences are color-coded to highlight the fluorescent protein regions, with eGFP in green, DsRed in red, and Epi-1 in pink, facilitating the schematic of the gene integration and orientation. The His-Tag of 6x histidine is highlighted blue and enterokinase site is highlighted yellow.

**Figure 2.**
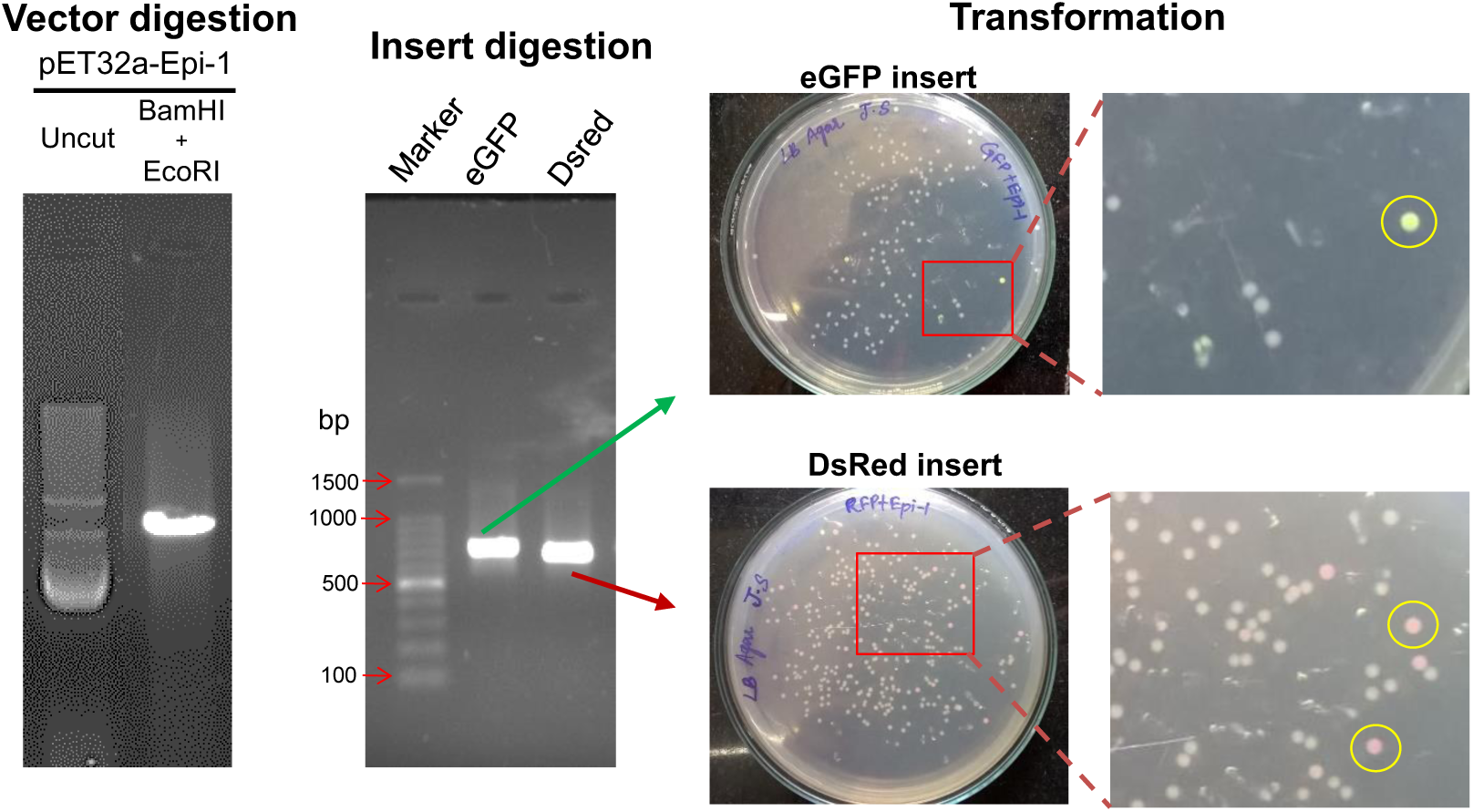
The image illustrates the molecular cloning steps involved in creating fluorescent Epi-1 gene clones. The experiment began with the digestion of the pET32a-Epi-1 vector using BamHI and EcoRI. Restriction digestion yielded a single linear band, while the uncut vector displayed supercoiled, nicked circular, and intact circular forms. The eGFP and DsRed inserts were PCR-amplified and digested with the same restriction enzyme pairs. The vector and fluorescent gene inserts were then ligated individually and transformed into competent cells *via* heat shock, followed by plating on fresh ampicillin-containing LB-Agar. After 48 hours of incubation, colonies that successfully took up the recombinant plasmids exhibited fluorescence corresponding to their respective inserts. Zoomed-in views show transformed colonies. The fluorescent colonies which are circled in yellow, indicate successful transformation.

**Figure 3.**
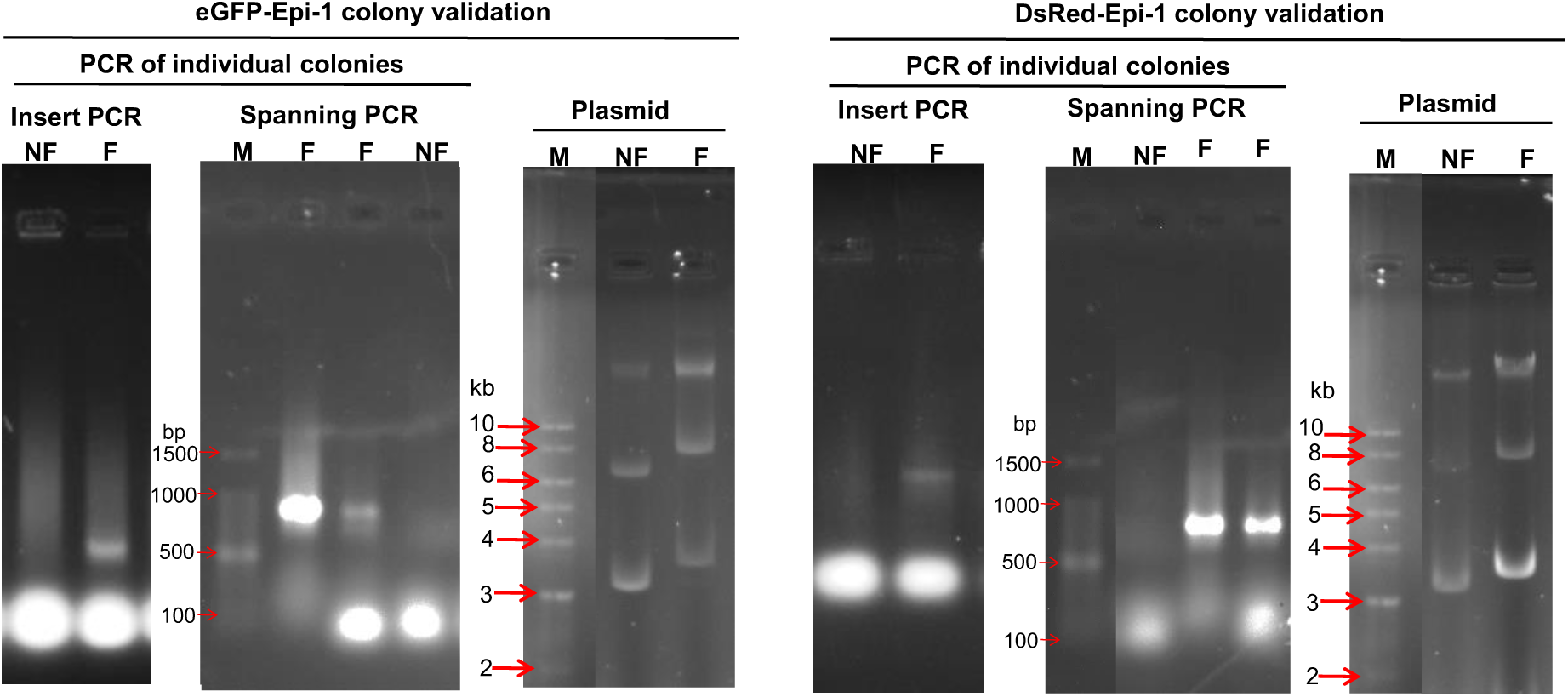
PCR and plasmid migration assay to validate the eGF-epi-1 and DsRed-epi-1 which were fluorescent screened in the previous figure. The figure shows agarose gel electrophoresis images of PCR (insert specific and spanning PCR) and plasmid isolated from individual clones of eGFP-Epi-1 and DsRed-Epi-1 and grouped individually. Individual **NF**-nonfluorescent colony, **F-**fluorescent colony are shown for representation. **M**-represents molecular weight DNA ladder. For insert PCR, the DNA isolated from fluorescent colony showed amplification with corresponding eGFP and DsRed primers. For spanning PCR, forward primer of fluorescent gene and and reverse primer of Epi-1 were used. The fluorescent colony showed amplification and the non-fluorescent colony did not show amplification. The size of the amplicon for eGFP-epi-1, is (717+84=801bp) and Ds-Red-Epi-1 is (685+84=769bp). The plasmids isolated from fluorescent (F) colonies migrate slower than the non-fluorescent (NF) colonies. This is because the non-fluorescent colonies has the same size of the vector (pET32a-Epi-1: 5944bp). The fluorescent colonies have a high number of bases due to insertion of fluorescent genes. eGFP-Epi-1: 661 bp and DsRed-Epi-1: 6629 bp.

### 3.3. Validation of visually fluorescent colonies by molecular techniques

To validate the visually screened fluorescent bacterial clones, molecular techniques such as PCR and plasmid migration assay were used. Insert specific and Spanning PCR resulted in fluorescent colonies containing bands in agarose gel, while the non-fluorescent colonies did not show any band in gel. Likewise the plasmid migration assay done for the fluorescent and non-fluorescent colonies reflected the result obtained from PCR. The fluorescent colony showed slower migration of plasmids compared to non-fluorescent colony. This is because the fluorescent gene insert caused a bigger plasmid size compared to vector alone.

## 4. Discussion

This study outlines the rationale and advantages behind designing fluorescent protein-based DNA constructs for the purpose of screening, fusion tag, overexpression and reporter assays for studying dynamic function of small antimicrobial peptides (AMPs).

The high cost associated with chemical synthesis and the low yields obtained from natural sources have prompted the exploration of recombinant bacterial expression systems as a more viable alternative for AMP production. Cationic AMPs pose a significant challenge in therapeutic development due to the need for large-scale production of highly purified compounds at competitive costs. Although chemical synthesis can produce both natural and modified cationic peptides, it is often prohibitively expensive and involves the use of hazardous reagents. In contrast, recombinant expression in host cells offers a more efficient and environmentally sustainable approach. Various recombinant systems have been explored for AMP production, including Escherichia coli, Staphylococcus aureus, insect cells, transgenic mammals, and plants [24]. Each of these systems presents unique advantages, and the success of anyone could enable scalable, cost-effective AMP production without the environmental burden of chemical synthesis. The key advantage of recombinant technology lies in its potential for long-term, sustainable peptide production—something that chemical synthesis cannot reliably offer.

AMPs and newly designed chimeric AMPs with multifunctions are produced in bacterial host by molecular cloning techniques. AMPs such as hepcidin, LL-37 [25], cecropin-melittin hybrid [26], indolicidin [27], serpin[28], crustin, lactoferricin [29] etc were produced by recombinant expression.

In general it is difficult to produce antimicrobial peptides in cassette vector because of their short peptide sequence, cationicity, unusual structure and high pI. In addition to the aforesaid factors the toxicity of small peptides [30-32] against the bacterial host cells and susceptibility to proteolytic degradation hinder the over expression of small peptides. These problems can be overcome by the expression of a peptide gene in fusion with a larger protein, followed by enzymatic or chemical cleavage to release active peptide [24]. The fusion partner serves to neutralize the positive charge of the peptide while providing some protection against proteolysis. In one study, Morin *et al*, [27] inserted a linker of short negatively charged spacer peptide sequence, that helped to neutralize the cations of the AMPs which acilitated the expression in the bacterial host. In the reviews by Li *et al*., [33-35] a comprehensive list of carrier proteins which were used for expression of AMPs are described. The carrier protein can be classified as solubility-enhancing carriers, aggregation-promoting carriers, self-cleavable carriers and secretion signal carrier proteins [36-38]. The soluble enhancing carriers can establish the fusion protein in soluble form in the host cytoplasm, example: Thioredoxin and glutathione transferase (GST). On the other hand the aggregation-promoting carrier proteins are reported to be more efficient than solubility-enhancing carrier proteins in protecting the host and masking the peptides from harming the host and segregating into inclusion bodies which helps in easy isolation and quick purification. PurF fragment, ketosteroidisomerase, PaP3.30, and TAF12 histone fold domain [33] are some of the examples of such category.

The goal of this study is to design a construct that would express a single fusion gene of GFP and AMP which can be over expressed as a fusion protein and used for localization of peptide within the live cells by confocal microscopy. In this case, the AMP is epinecidin-1 is fused with eGFP and DsRed at its N-terminus.

GFP has been reported as a more stable carrier protein for expressing AMPs [39] and it has unique property of allowing its detection within the live cells. In one study, GFP fused small AMP peptide such as tachyplesin partitions into inclusion bodies facilitating easy purification [40] and it helps in extraction of small polypeptides with physiological buffers.

For the purpose of screening, the bacterial transformants carrying a recombinant plasmid with the gene of interest has become more rapid and simple because it can be visually detected. GFP as a fluorescent probe lacks the requirement for an exogenous cofactor [41]. GFP can be expressed in intact tissues and processes can be monitored without the disturbance caused by the introduction of reagents. This saves time and money, more importantly, less effort and manpower will be neededto screen the correct clones. In this study, we cross verified the fluorescent colonies with gene specific and spanning PCR compared to the non-fluorescent colonies. The fluorescent colonies had the insert in correct orientation.

As a biological marker, GFP or DsRed expression are visually observable when expressed in bacteria. The use of GFP as a tag does not alter the normal function or localization of the peptide or protein [42]. GFP has found its broad use in almost all organisms and all major cellular compartments. In cellular biology, GFP has been used as reporter gene, cell marker, fusion tag, indicator for protease action, calcium sensitizer [43] [44]etc.

GFP can also be called as a tracker molecule since it has been used in intracellular trafficking such as migration dynamics of proteins within subcellular sites, sensors of neuronal membrane potential etc. It is easy to find out where GFP is at any given time by shining ultraviolet light. For instance, GFP attached to a the AMPS, can be actively monitored through green glow inside the host (Helps in answering questions such as does it reach to the cytoplasm or the membrane or any specific organelle?).

GFP cloned in an expression vector can be screened for positive clones with the luminescence observed with the naked eye. The intensity of fluorescence would also indicate the extent of solubility of the fusion protein. Banerjee et al., [45] have shown that the solubility of the target protein could be predicted in situ at the time of recombinant screening based on the intensity of the GFP fusion proteins. They demonstrated that higher the solubility of the target protein, the intensity of the GFP fluorescence on the agar plate is higher rendering the screening of the recombinants a dual objective of identification and predicting the solubility of the gene of interest attached to the reporter gene. If the fusion protein is expressed as insoluble aggregate, the *E. coli* has shown less or no luminescence [46]. The soluble *E. coli* protein exhibits higher intensity of luminescence or fluorescence. Thus with this one molecule multiple functionalities can be studied.

Embarking the functionalities of GFP or DsRed as a carrier protein we have successfully cloned the fluporescent eGFP gene and DsRed gene in frame with epinecidin-1 by recombinant molecular biology technique and ascertained the presence of gene in-frame. These clones shall be proceeded with over expression and purification. One of the challenge associated with the expression is the expression of AMPs in soluble form. At the initial time points, as the protein is expressed in lower quantities a bright fluorescence is seen, However when the plates were incubated for a long time, a drop in fluorescent signal was observed which may be a sign of cell death due to higher expression of antimicrobial peptide. This will be a drawback when aiming for expression in large bioreactors. Hence continuous batch collection harvest will be needed to be devised for oprimum harvest of desired protein.

## 5. Conclusion

In this study, eGFP-Epi-1 and DsRed-Epi-1 were successfullyconstructed using molecular cloning methods and confirmed using PCR and plasmid migration assay.

## 6. Acknowledgement

We thank Indian Council of Medical Research, Govt. of India, New Delhi for partial financial support *via* sanctioned research project to the corresponding author (Ref. No. IIPSG-2024-01-01898).

## 7. Author Contributions

Conceptualization: S.J and A.K.; methodology: S.J, J.J and A.K.; validation, S.J., J.J. and A.K.; formal analysis, S.J, J.J and A.K.; investigation, A.K.; resources, S.J., and A.K.,; data curation, S.J., J.J and A.K; writing—original draft preparation, S.J. J.J., H.R., and A.K.; writing—review and editing, S.J. J.J, H.R, and A.K.; visualization, S.J. and A.K.; supervision, A.K.; project administration, A.K.,; funding acquisition, S.J., A.K. All authors have read and agreed to the published version of the manuscript.

## Notes

### Competing Interest Statement

The authors have declared no competing interest.

